# Linking Engineered Cells to Their Digital Twins: a Version Control System for Strain Engineering

**DOI:** 10.1101/786111

**Authors:** Jonathan Tellechea Luzardo, Charles Winterhalter, Pawel Widera, Jerzy Kozyra, Victor de Lorenzo, Natalio Krasnogor

## Abstract

As DNA sequencing and synthesis become cheaper and more easily accessible, the scale and complexity of biological engineering projects is set to grow. Yet, although there is an accelerating convergence between biotechnology and computing science, a deficit in software and laboratory techniques diminishes the ability to make biotechnology more agile, reproducible and transparent while, at the same time, limiting the security and safety of synthetic biology constructs. To partially address some of these problems, this paper presents an approach for physically linking engineered cells to their digital footprint - we called it digital twinning. This enables the tracking of the entire engineering history of a cell line in a specialised version control system for collaborative strain engineering via simple barcoding protocols.

## 2. Introduction

There is a rapidly accelerating convergence between biotechnology and information and communication technologies (ICT).

As writing and reading DNA becomes routine, cheaper and pervasive, the genetic engineering – that is, the programming of biological organisms – becomes more similar to programming computers. This has important disruptive implications for biotechnology: (1) larger teams of bio-programmers – both in biotechnology companies of all sizes and in research organizations such as universities – work together and concurrently in the genetic programming of biological organisms; (2) the pace of innovation for biological “apps” is set to explode and (3) the ecosystem and supply chain of biotechnological products, in particular cell lines and plasmids, is to become more complex and more diversified.

In software development, the time to market for new products has shortened dramatically over the recent years, thanks to (amongst other factors) advances in *continuous integration and deployment* allowing teams of programmers (often geographically distributed), to rapidly develop, test, debug and promptly push new code to production. This means that new software products can reach their users and customers faster than ever before.

The core element of the technology underpinning continuous integration and deployment are the Version Control Systems (VCS). Their roots date back to Bell Labs’ early 1970’s Source Code Control System (Rochkind, 1975). Git, the prevalent VCS powering modern software engineering, was introduced by Linus Torvalds in 2005 and popularised the use of *distributed* version control systems.

Nowadays distributed VCS are used daily by software developers across the world. VCS help distributed teams to keep track of the *history of changes* to large software systems as it enables them to quickly answer questions such as *“What was done to a piece of software?”, “Who introduced the last feature (or bug) into the code?”, “When did the modification take place?”, “What was modified?”, “How does this new version differ from the previous one?”, “What versions of the code are available?”*, etc.

A version control system provides answers to the above questions, greatly simplifying the work of distributed teams of programmers who must modify the same piece of software concurrently. VCS achieve this, by storing the files in the repository together with the entire history of changes to them; changes, are automatically tracked and semi-automatically merged by the system. Because the entire *lineage* of a source code is maintained, VCS enables traceability, transparency and backtracking (if necessary) as well as branching out new versions of computer code. Branched code does not interfere with the original code and yet retains all the metadata that lead to that branch’s origins, thus providing a safe sandpit for experimentation (i.e. if something breaks, one can always restore previous states). Without VCSs it would be virtually impossible to develop complex computer programs as we know them today. For a recent biology and biotechnology friendly introduction to version control and Git refer to (Blischak et al., 2016).

How would a biotechnology-specific version control system contribute to making biotechnology more agile, reproducible and transparent? —and the corresponding live constructs altogether tractable for the sake of their security and safety? (Schmidt et al., 2016)

Mislabelling and misidentification of biological samples occurs frequently in the laboratory (Broman et al., 2015). While being able to properly identify and track a strain’s origins is essential, and notwithstanding persistent calls to address this challenge (American Type Culture Collection Standards Development Organization and Workgroup ASN-0002, 2010; “Identity crisis,” 2009; Masters, 2012), it is a problem that lacks appropriate tooling. Moreover, a related major problem affecting many scientific disciplines - including biotechnology- is experimental irreproducibility (Freedman et al., 2015). As recently as 2018 Nature had a special issue themed *“Challenges in Irreproducible Research”* to highlight some of the most urgent issues and suggest potential ways forward.

Laboratory data generated by researchers also face problems like data loss and lack of accepted engineering and reporting standards. Electronic Lab Notebooks and other online applications that use the cloud to store and publish the data and experiments generated are a partial step in this direction although they are generic for lab operations rather than specific to the genetic programming of organisms. Thus new tools and software tools have been called for (Sadowski et al., 2016).

Moreover, although several methods aimed at identifying cell lines coming from clinical or environmental samples have been developed, little effort has been put into establishing such tools for laboratory-created strains. From an engineering view point, one needs to know not only a strain’s genome sequence but, importantly, also the history of changes made it to (e.g. added plasmids, scars left by knock-ins/knock-outs…), experimental conditions used, the intention behind the modifications and other metadata such as, e.g., the lab of provenance, the genetic engineer(s) who worked on a strain, etc.

The combination of the issues mentioned above lead to the wasteful use of both public and private time and money, to poor biotechnology practices as well as to public mistrust due to the lack of trackability and transparency in biotechnological product innovation.

A further crucial point that has gone unnoticed but that has important implications is that a strain’ digital footprint (e.g. strain designs, genome sequence, recombineering sites, engineering history, etc) is *disconnected* from the physical strain sample. That is, no actual connection exists between the strains one creates in the laboratory or that is used in production and the data available for that strain. Current practice for linking a strain digital footprint to the actual biological sample is based around hand-written notes, word processor or spreadsheet files and – in the best case – the labelling of test tubes with barcodes generated from generic laboratory information management systems. Crucially, in all those cases, the biological sample itself does not carry a record of its digital footprint.

Recent advances in DNA synthesis techniques and information storage in DNA could help bridge this important gap. The usage of DNA as a long-term way of storing data has been recently proven and exploited (Shipman et al., 2017). The use of small, synthetic DNA sequences as identifiers has been reported several times in very diverse areas. These short DNA sequences allow the identification of mutants in mixed populations in both microbial studies (Liu et al., 2017; Mazurkiewicz et al., 2006) and tumour cell lines studies under different treatments (Bhang et al., 2015; Yu et al., 2016). DNA barcodes can be used in gene synthesis, in which the barcode sequence is used to isolate and assemble the synthetic gene (Plesa et al., 2018). Additionally, DNA barcodes have been shown to be useful in drug discovery by tagging and identifying several chemicals that bind to specific target molecules (Zimmermann and Neri, 2016). Very recently new barcode generation algorithms have been developed to consider any type of synthesis or sequencing mistake which increases the robustness of all the previous uses (Hawkins et al., 2018).

Finally, regardless of the many proposals of genetic firewalls for containing genetically engineered organisms and SynBio agents, reality is that current metrics (see above) never go beyond events occurring at frequencies of 10-11, what is not enough for what has been called *certainty of containment* (CoC) (Schmidt et al., 2012). There is widespread opinion in the Life Sciences community that no firewall, sophisticated as it might be, will stop engineered organisms to scape a given niche (de Lorenzo et al., 2018). While CoC is a fascinating scientific question, most of the concerns on undesirable propagation of human-made constructs can be managed if chassis and agents were barcoded with specific and unique DNA sequences which— once decoded—could take users to the most detailed information available for this or that particular construct. Barcoded clones would thus be equivalent to pets implanted subcutaneously with identification chips: in case they get lost or do some harm, their owner and their pedigree can be immediately identified. By the same token, *barcoded* strains would allow accessing all relevant information on its pedigree, safety and modifications implemented in them. This will be ultimately more useful that any containment measures—which in all cases are bound to fail. Instead, barcodes will not only make traceability simple, but it will also assign a non-ambiguous cipher to the growingly improved versions of the same chassis (as is the case with computers and mobile phones operating systems).

In this paper we present a biotechnology specific version control system, CellRepo, that provides both the genetic toolkits and cloud-based software to physically link living samples to their digital footprint history. CellRepo is based on small, unique and bio-orthogonal DNA sequences inserted in specific genomic locations of a strain. By a single sequencing reaction, the DNA sequence can be retrieved, and hence the strain user can track down the entire digital footprint history of a strain via our web server: strain creators, parental and derivative strains, strain design documentation, related papers, experimental protocols, computer models, etc can all be retrieved via the cloud computing component of our system (Fig. 1). Put together, the biotechnology kits and the software repository move the digitalization of biotechnology a step closer making it more collaborative, scalable, transparent, trackable and reproducible.

**Figure 1:**
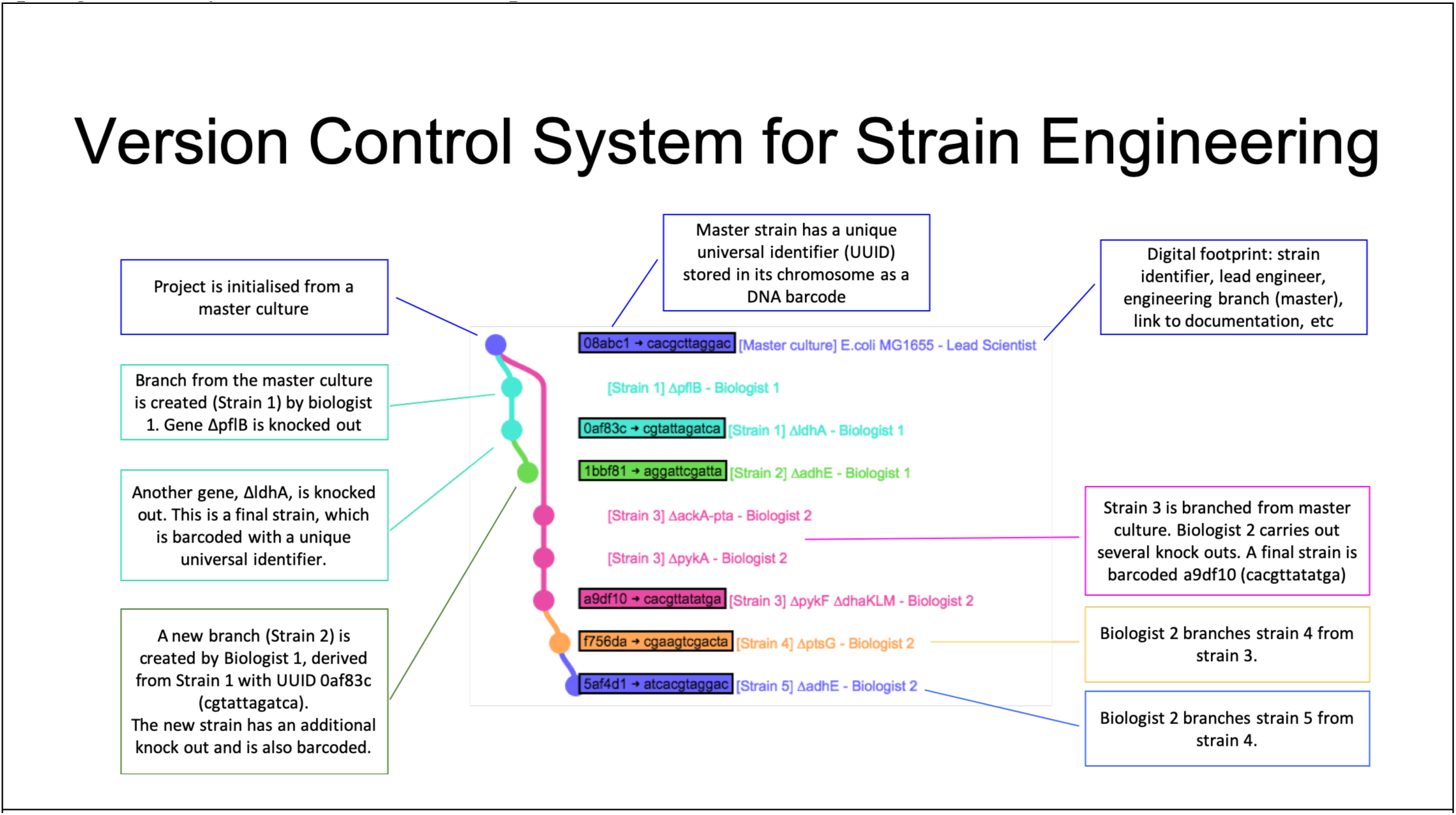
An example of version control for strain engineering. A master strain repository is created (blue) and from there two main branched strain projects are derived (cyan and pink). Each “commit” (coloured dot) to the repository represents a key strain engineering milestone. The commit has a unique identifier, the DNA barcode, that is inserted into the chromosome of the strain.

## 3. Materials and methods

We report next the materials and methods used for physically relating a given strain to its digital footprint within the cloud-based version control system. This is accomplished via writing and reading of a unique DNA barcode (Fig. 2) as explained next.

**Figure 2:**
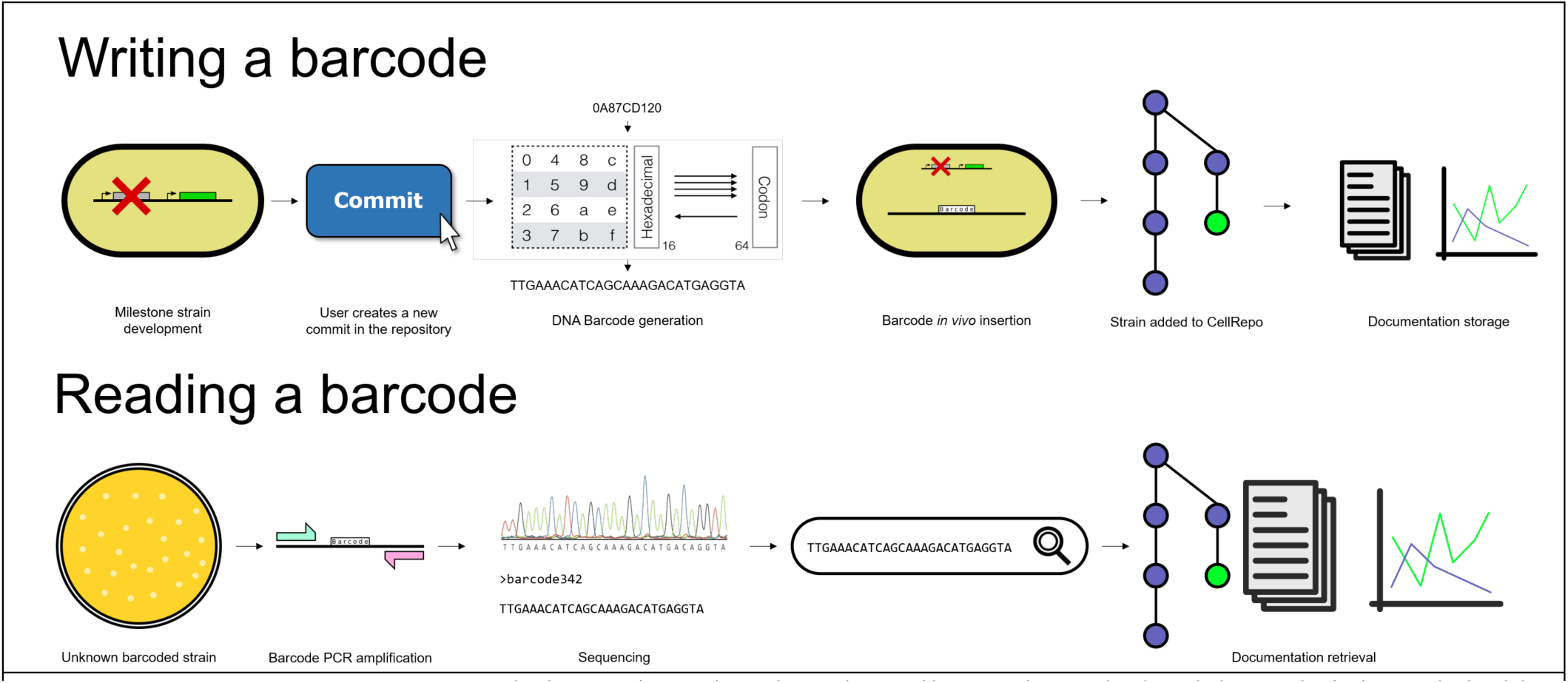
**Writing a barcode (Top) –** during strain engineering a key milestone is reached and the strain is barcoded with a unique DNA sequence derived from CellRepo cloud-based repository. A commit to the repository is made connecting the unique barcode (now in the chromosome of the cell) with all the cell’s related documentation. **Reading a barcode (Bottom) -** a DNA barcode is read from a sample strain enabling the lookup of all the strains’ documentation from the CellRepo cloud-based repository.

### 3.1 Materials

DNA was amplified using Q5 High-Fidelity DNA Polymerase (New England Biolabs, NEB). For cloning purposes DNA was purified using Monarch PCR & DNA Cleanup Kit (NEB) and assembled using NEBuilder HiFi DNA Assembly Cloning Kit (NEB). Plasmid preparations were carried out using QIAprep Spin Miniprep Kit (QIAGEN). Primers and Synthetic DNA sequences (barcodes gBlocks) were synthesized by Integrated DNA Technologies.

### 3.2 Strains and plasmids

*E. coli* DH5-α cells were used for most of plasmid preparations. Plasmids carrying a R6K-ɣ origin of replication were prepared and stored using DH5-α/λ-pir cells. BW25113 strain was used as a barcode receiver and as genomic DNA template source for homologous regions cloning for *E. coli* experiments. For *B. subtilis* experiments, 168 strain was used as a barcode receiver and as genomic DNA template source for homologous regions, strain ZPM6 was used for toxin/antitoxin experiment (Lin et al., 2013).

All the “vector” tagged plasmids were built containing a restriction site that allows easy barcode sequence cloning. See Table 1 for a full list and description of the plasmids utilized in this work.

**Table 1:**
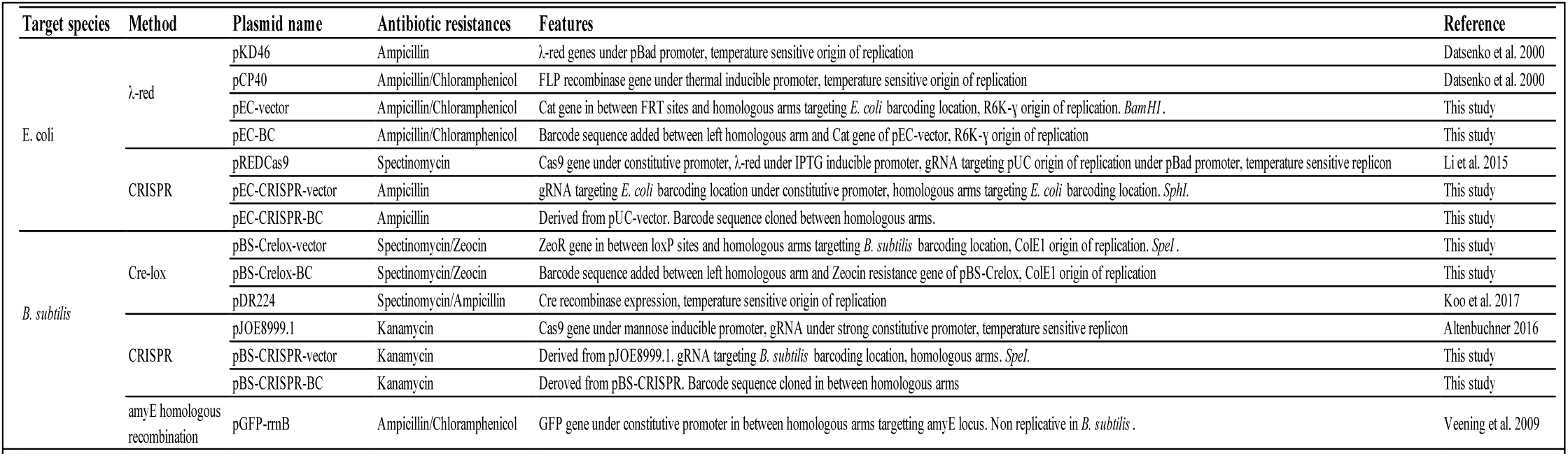
Plasmids used in our barcoding kits.

### 3.3 Barcoding site selection

The barcodes that irrevocably link a strain to its digital twin must be stably inserted in the chromosome in a manner that is both stable and does not change a strain’s phenotype. For this reason, we choose to insert the barcodes far from important gene loci. On the chromosome, interacting regulatory units are often found in neighbouring locations. Not all genes in the genome are essential and therefore, for each species, we gathered and curated data about essential genes from the literature and ruled out locations in the genome that were neighbouring any essential units. Genes involved in metabolism regulation, cell wall components and migrating elements were also avoided. Additionally, the barcoding loci had to be conserved between *E. coli* or *B. subtilis* lab strains. Following these considerations, barcodes were placed at loci that, upon cell division, replicates properly but -at the same time-have a favourable biological context, namely, the barcodes at these loci (fig. 3) are unlikely to interact with proximal elements or to interfere with strain specific genetic circuitry.

**Figure 3:**
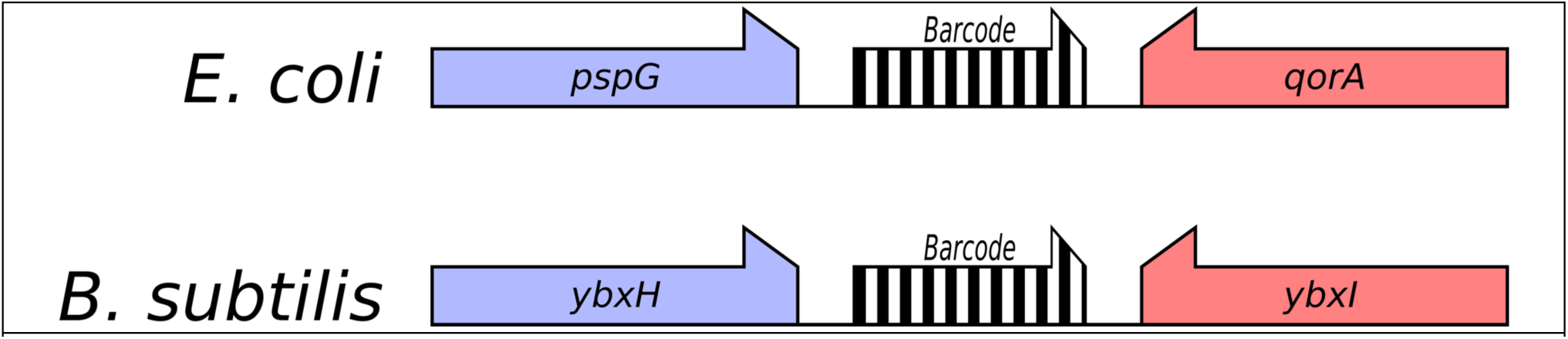
Graphic representation of the barcoding sites for both species. For both species a non-essential gene pair was chosen.

### 3.4. *E. coli* barcoding

#### 3.4.1. λ-Red recombineering

pEC-vector was assembled containing the R6k-γ origin of replication and a selection cassette (*cat* gene, conferring chloramphenicol resistance) flanked by FRT sequences and the homologous arms. Once the vector plasmid was ready, the barcode sequence was cloned into it. The whole process followed an adaptation of the protocol described by (Datsenko et al., 2000). Competent cells were prepared, transformed with plasmid pKD46 and grown at 30°C in LB agar supplemented with carbenicillin (100 µg/mL). Transformant cells were induced with L-arabinose 30 mM.

Barcoding cassette was amplified by PCR from pEC-BC. Electrocompetent cells were prepared and mixed with the barcoding cassette. Cells were zapped and plated in LB plates supplemented with chloramphenicol (25 µg/mL) at 37°C. Barcoding was checked by colony-PCR. If selection cassette removal was required, barcoded clones were transformed with pCP20 plasmid and incubated at 30°C in LB/Carbenicillin plates. Carbenicillin resistant colonies were grown at 37°C in LB plates. Carbenicillin and chloramphenicol sensitive clones were checked by colony PCR. Positive clones were stored as barcoded strains.

#### 3.4.2. CRISPR

*E. coli* cells were barcoded by CRISPR using a two-plasmid system (Jiang et al., 2015; Li et al., 2015). The plasmid pEC-CRISPR-vector was constructed by cloning the homologous arms and the 20N-gRNA scaffold under a constitutive promoter. The barcode sequence was added afterwards in between both homologous arms by HiFi DNA Assembly. pREDCas9 was transformed into BW25113 cells and selected in LB/Spectinomycin (50 µg/mL). λ-Red recombinase was induced with IPTG 2 mM until OD=0.5. pEC-CRISPR-BC was transformed and selected in LB/Spectinomycin/Carbenicillin plates. Transformant cells were checked by colony PCR. Positive clones were cured from pEC-CRISPR-BC by L-arabinose induction (30 mM). pREDCas9 was cured afterwards by growing cells at 37°C. Spectinomycin and carbenicillin sensitive clones were stored.

pREDCas9 was a gift from Tao Chen (Addgene plasmid # 71541).

### 3.5. *B. subtilis* barcoding

#### 3.5.1. Toxin/Antitoxin

Using SOE-PCR a cassette containing both homologous arms, a mazF-ZeoR cassette and the barcode sequence was created and amplified following an adaptation of the protocol described in (Lin et al., 2013). After transformation colonies were restreaked on LB/Zeocin (20 µg/mL) plates and tested for the integration of the recombinant DNA by PCR. A positive clone was grown with xylose (1%) and the toxin gene induced. Cells were plated in xylose supplemented media. Individual colonies were restreaked on LB and LB/zeocin plates and colonies were tested positive by PCR and sequencing.

#### 3.5.2. Cre-Lox

An adaptation of (Koo et al., 2017) protocol was used. A vector containing the antibiotic resistance gene (Zeocin) flanked by loxP sites and homologous arms was created. Barcode sequence was added afterwards. Barcoding cassette was amplified by PCR and transformed into 168 cells. Zeocin resistant cells were transformed with pDR244 at 30°C. Spectinomycin (100 µg/mL) resistant cells were checked by colony PCR. pDR44 was cured at 37°C. Spectinomycin/Zeocin sensitive cells were stored.

#### 3.5.3. CRISPR

To barcode *B. subtilis* cells using CRISPR we used a single-plasmid approach (Altenbuchner, 2016). Homologous arms and sgRNA target sequence were cloned in pJOE backbone. The barcode DNA sequence was cloned in afterwards. 168 cells were transformed with this plasmid and selected in LB/Kanamycin (5 µg/mL) supplemented with 0.2% mannose for Cas9 induction at 30°C. Transformants were checked by colony PCR. pJOE was cured by growing the cells at 37°C. Kanamycin sensitive cells were stored.

### 3.6. Checking barcode sequence and presence

To easily check the integrity of the barcode sequences, the identifier was tagged with a Universal Primer sequence (TGGACATACATAGTATACTCTGGTG). This primer is used in the Sanger sequencing reaction to check the sequence of the barcode. Also it can be used to check the success of the barcoding experiment (through colony-PCR) together with appropriate species-specific reverse primers.

### 3.7. Barcode stability assay

#### 3.7.1. Chemostat

Using a chemostat we followed bacteria over 200 generations (about 4 days of culture) and retrieved barcode information at regular time points (every 15-25 generations). For *E. coli* the same cultures were followed over 200 consecutive generations, while in *B. subtilis*, cells were followed for 100 generations, induced to stress and sporulation by ethanol treatment and regrown from spores for a further 100 generations. Barcode sequences were obtained by PCR product Sanger sequencing after barcode amplification from genomic DNA.

For this study, we performed CFU growth curves and serially diluted cultures over time to obtain ideal dilution rates at which a minimal number of cells could be used to inoculate a chemostat. Provided this minimum number of cells, it was possible to evaluate, given a specific growth rate, the time needed for cultures to attain exponential phase (e.g. when the chemostat continuous flow should be turned on). Along all chemostat experiments, we recorded optical density (OD) measurements while sampling cells for barcode sequencing to ensure estimated dilution rates were accurate and cultures could reach steady-state growth.

#### 3.7.2. Large-scale growth assay

With automated plate handling systems, we compared the evolution of 384 subcultures of barcoded and control cells over several stationary phase redilutions, in order to observe any changes represented in growth defects. Besides growth characterisation, all barcoded samples were sequenced to uncover any potential mutations in DNA barcode sequences.

We used a liquid handling robot (Beckman Coulter Biomek FX) and an automated plate reader to handle 8 individual 96-well plates simultaneously. Before starting a growth experiment, plates were sealed by a gas-permeable membrane. In this assay, we followed bacteria over 10 subculture experiments. Initially, a single colony from each barcoded/control strain was picked from a fresh plate and grown in 25ml LB supplemented with 0.4% (w/v) glucose overnight at 37°C with regular shaking parameters (about 150 rpm). In the morning, saturated cultures were spun down, resuspended in fresh LB medium, diluted 100 times and 200µl were loaded onto ThermoFischer clear 96-well microplates.

For all subculture experiments, two conditions were tested: an early stop of bacterial cultures after 6h (in late exponential/start of stationary phase) and a prolonged culture in stationary phase (12h) before snap freezing. For the first subculture, 100µl were harvested after 6h for the early sampling point, and the remaining bacterial culture was further incubated up to 12h. All subsequent cultures were diluted 100 times from frozen stocks and cultivated in a 100 µl total volume for both early and late sampling points. By the end of the 10 subcultures, genomic DNA (gDNA) was extracted and screened for potential variations in barcode sequences.

## 4. Results and Discussion

We illustrate all the concepts introduced by performing a simple genetic engineering experiment that includes barcoding a cell line and uploading to the version control system. *B. subtilis* 168 wild-type strain was barcoded with Barcode 659 using the Cre-lox method described before. The resulting strain was then transformed with pGFP-rrnB (Veening et al., 2009). Chloramphenicol (5 µg/mL) resistant clones were checked by colony-PCR and by checking the green fluorescence emission in a plate reader. This new strain was re-barcoded using Barcode 207.

### 4.1. Barcoding process

All the methods used to barcode both species probed to be capable of barcoding the cells with a high efficiency (>90%). After extracting the genomic DNA and PCR the barcode, it was always possible to retrieve the barcode DNA sequence.

### 4.2. Barcode stability assay

Using the chemostat we analysed 128 sequencing reactions after 200 generations, including 24 controls to compare the evolution of barcoded vs. wild-type strains. We confirmed that control wild type sequences remained unchanged and found no variation in barcode sequences over 200 generations for either species.

In the large-scale growth assay, over 10 subcultures, we estimated from 100-fold dilutions of previous subcultures that final samples reached about 100 generations. In our assay, sequencing of 384 barcoded strains tested in a normal vs. stress conditions always revealed intact DNA barcode sequences. No major difference in growth rates was observed across the different samples. The majority of sequencing reads left a 26-27 nucleotide gap downstream of the universal primer-binding site and then showed a perfect match with the expected alignment. In less than 5% of cases, sequencing data quality was noisy, but a second complementary read would always manage to recover the integrity of a barcode sequence. The screen of a large number of biological replicates helped us to assess the robustness of barcode sequence insertion in the bacterial genome and demonstrated the stability over time of the barcodes.

### 4.3. Web server

We implemented the idea of CellRepo as a web application. We used CellRepo to document the key milestones throughout the process (fig. 4) of cell engineering, characterisation and barcoding described earlier.

**Figure 4:**
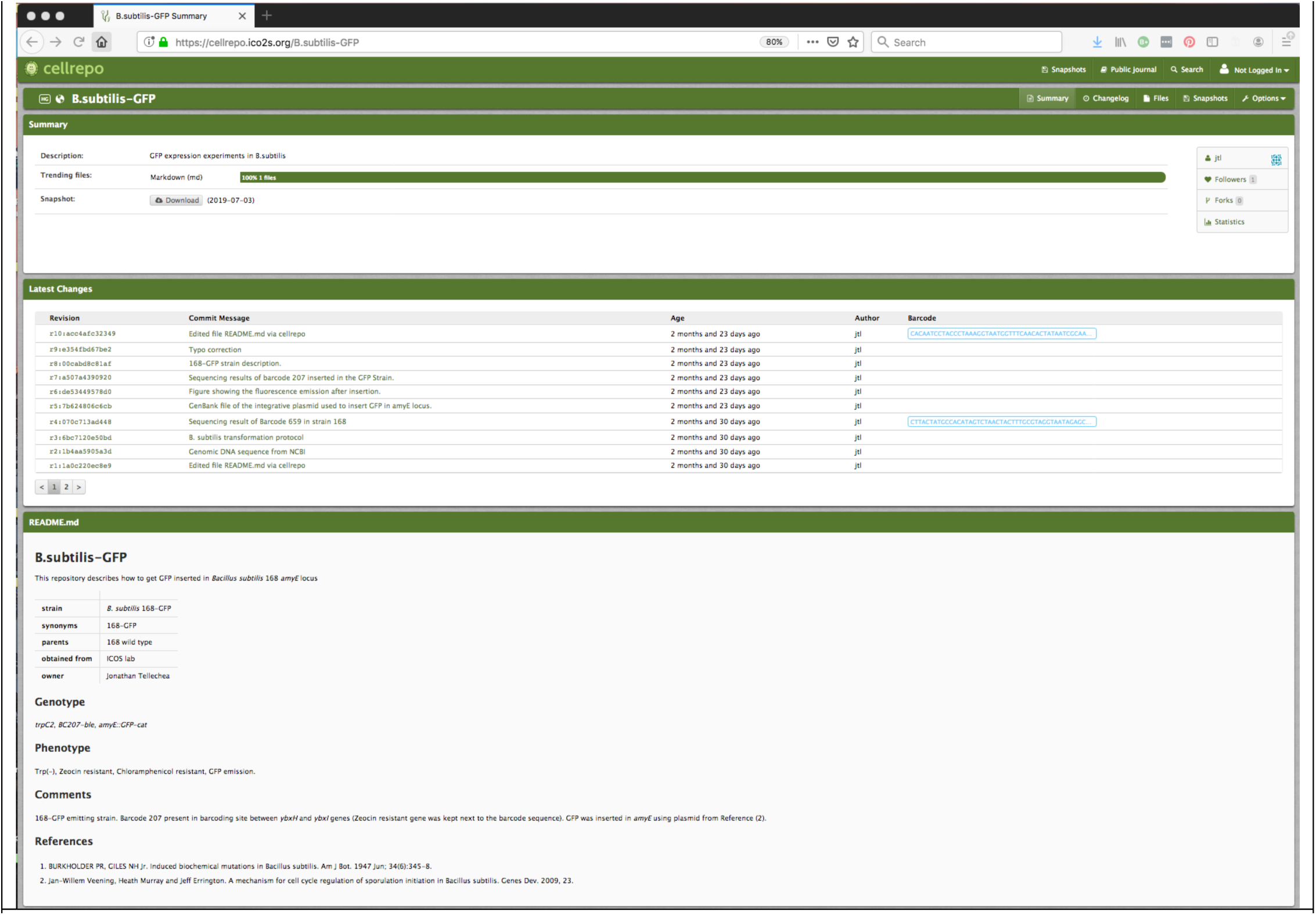
Overview of the cell repository that includes the various “commits” during cell engineering of a bacillus subtilis mutant that includes a GFP gene. All the revisions to the digital footprint of the cell line are visible with two key engineered milestones linked via a genetic barcode to the cell line.

The web application, which is available at https://cellrepo.ico2s.org, can be freely used. It provides functionality to register a new user, who can then proceed to create a number of different cell repositories (cellrepos). The owner of a cellrepo can commit -i.e. store-data to it and generate barcodes. There is no limit on the type of data that can be associated with a cell engineering repository. Data might include plasmid designs, FASTA or GenBank files with genomic or plasmid data, SBOL files, characterisation experiments outputs (e.g. optical density readouts, growth curves, experimental protocols, etc.) references to papers or the papers themselves, etc. All commits are organised in chronological order. Repositories have a wiki-like front-page that describes the essential details (e.g. genotype, phenotype, owner, etc.) of the project. Furthermore, the system allows a repository to be forked so further work could be carried into a cell line without interfering with the original repository (fig. 5).

**Figure 5:**
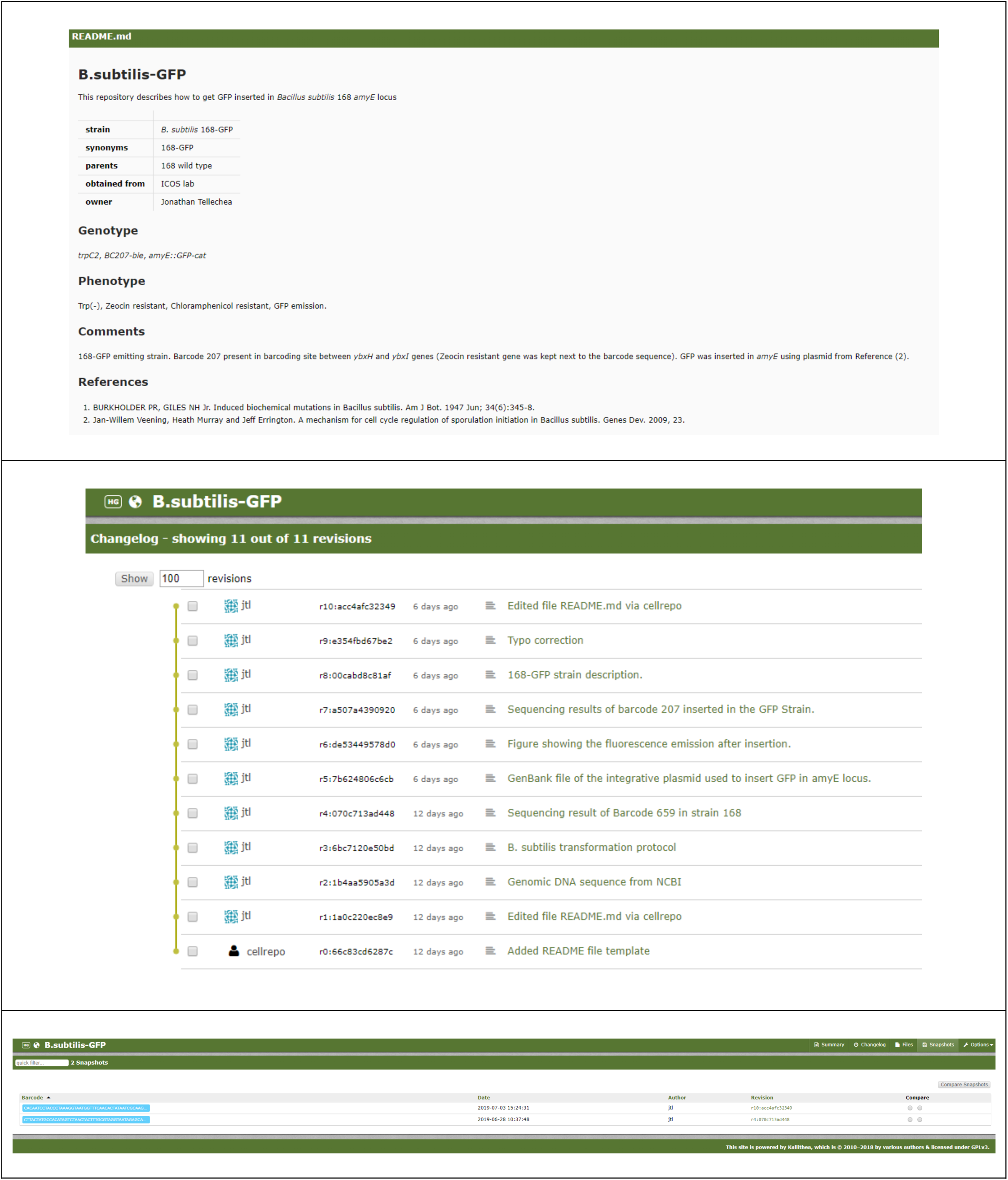
(top) Essential cell engineering repository information. **(middle)** List of cell engineering commits. Commits can be expanded to show detailed information and associated files. **(bottom)** Some of the commits (in this example those with revision id r10:acc4afc32349 and r4:070c713ad448) have been barcoded back into the cell line. These commits are called “snapshots” as they uniquely link the cell line to the snapshot of all the digital documentation at that point in the cell engineering cycle.

## 5. Conclusions

In this paper we argue that, as the speed, size and complexity of synthetic biology, biotechnology and genetic engineering approaches increase, new tools are required to handle the substantial scale up that is taking place. We propose a new purpose-built version control system, CellRepo, for strain engineering. CellRepo links biological cells to their “digital twins” thus allowing the tracking of all extant data and metadata related to a cell engineering process. This link is created by placing a barcode into the cell that can -at a later stage-be retrieved via a simple sequencing reaction. The retrieved barcode can then be used to identify the cell line and all the related details stored in CellRepo (who built the cell line, in which lab, when the modifications took place, what protocols were used, etc).

As the need of barcoding new species and Synthetic Biology chassis increases, new genetic technologies need to be developed for stably inserting such sequences in the genome of target organisms though all the taxonomic scale. In this regard, we envision that adoption of barcoding as a routine for standardized identification of genetically engineered organisms will ease approval, security and safety of the corresponding modified agents for industrial and environmental uses to a degree far superior than the current propositions for genetic firewalls—which by no means provide a certainty of containment. Besides the computational effort, such regulation-oriented and safety-oriented barcoding will demand genome editing of strains deficient in recombination, a feature typically requested for environmental safety of GMOs. To this end, a number of molecular tools e.g. counterselectable TargeTrons are being currently developed in our Laboratories (Velázquez et al., 2019). In the meantime, CellRepo is freely available to use at https://cellrepo.ico2s.org

## 6. Acknowledgements

JTL, PW, JK, CW and NK were supported by the UK Engineering and Physical Research Council under project *“Synthetic Portabolomics: Leading the way at the crossroads of the Digital and the Bio Economies (EP/N031962/1)”*. VdL was supported by project *“BioRoboost (H2020-NMBP-BIO-CSA-2018, grant agreement N820699)”*.

